# Mechanical forces in extendable tissue matrix orient cell divisions via microtubule stabilization in *Arabidopsis*

**DOI:** 10.1101/2023.01.16.524206

**Authors:** Lukas Hoermayer, Juan Carlos Montesinos, Leonhard Spona, Saiko Yoshida, Petra Marhava, Silvia Caballero-Mancebo, Eva Benková, Carl-Philip Heisenberg, Jiří Friml

**Author notes:** Corresponding author: Jiří Friml.

## Abstract

Plant morphogenesis relies exclusively on oriented cell expansion and division. Nonetheless, the mechanism(s) determining division plane orientation remain elusive. Here we studied tissue healing after laser-assisted wounding in roots and uncovered how mechanical forces of cell expansion stabilize and reorient microtubule cytoskeleton for orientation of cell division. We revealed that root tissue functions as interconnected cell matrix with a radial gradient of tissue extendibility causing a predictable tissue deformation after wounding. This causes instant redirection of expansion in the surrounding cells and reorientation of microtubule arrays ultimately predicting cell division orientation. Microtubules are destabilized under low tension, whereas stretching of cells, either through wounding or external aspiration immediately induce their polymerisation. The higher microtubule abundance in the stretched cell parts leads to reorientation microtubule arrays and ultimately cell division planes. This provides a long-sought mechanism for flexible re-arrangement of cell divisions by mechanical forces for tissue reconstruction and plant architecture.

## Introduction

Cells in plants are fixed in their position within a tissue; they are surrounded by a cell wall matrix that prevents cell migration, limits their expansion and glues them together. Owing to this immobility, oriented cell division and expansion are the sole determinants of morphogenesis. The division plane orientation defines how two daughter cells will be separated spatially by a new cell wall; hence, is crucial for cell type acquisition and tissue development (Dolan *et al.*, 1993a; van den Berg *et al.*, 1995; Rasmussen and Bellinger, 2018).

A crucial developmental process relying on precisely executed and directed cell divisions is wound healing, where damaged patches of cells are replaced by proliferating cells (Bloch, 1941; Hoermayer and Friml, 2019; Ikeuchi *et al.*, 2019). Recently, we introduced an experimental setup of cell ablation in the root meristem that allows the investigation of tissue restoration on a cellular level, *in situ* and in real time (Marhava *et al.*, 2019; Hoermayer *et al.*, 2020). Our studies revealed the phenomenon of restorative divisions, where cells switch division planes to invade and replace removed cells through subsequent cell expansion and cell fate acquisition. These restorative divisions occur predominately at the inner adjacent side of the wound and show similarities to the regeneration of the whole-root excisions (Xu *et al.*, 2006; Sena *et al.*, 2009; Efroni *et al.*, 2016; Marhava *et al.*, 2019; Hoermayer *et al.*, 2020). However, the underlying mechanism behind those precisely executed cell divisions and their exclusive occurrence in inner adjacent cells remain elusive.

How cell division planes are placed in root tissues is crucial for regulating tissue architecture (Dolan *et al.*, 1993b; van den Berg *et al.*, 1995; De Rybel *et al.*, 2013, 2015) and outgrowth of new organs like the lateral root (Dubrovsky *et al.*, 2008; Marhavý *et al.*, 2016; Vilches Barro *et al.*, 2019). Hence the division plane selection is highly regulated and multiple influencing factors have been described: geometry, developmental cues, local and tissue-derived stress (Rasmussen and Bellinger, 2018). The default choice for cell division plane is determined by the cell geometry: a plant cell prefers division perpendicular to its longest axis, with the smallest possible new cell wall area (Errera, 1888; Besson and Dumais, 2011). However, this rule does not explain the orientation of divisions in most plant cells. Developmental cues may override the division plane selection through gene expression or localisation of polarity-defining proteins, e.g. during formative divisions in stem cells and throughout embryo development (Helariutta *et al.*, 2000; Heidstra, Welch and Scheres, 2004;

Willemsen *et al.*, 2008; van Damme *et al.*, 2011; De Rybel *et al.*, 2013; Yoshida *et al.*, 2014; Rodriguez-Furlan *et al.*, 2022). However, such developmental cues only mildly affect the division plane switch after cell ablation (Marhava *et al.*, 2019), suggesting a greater importance of alternative factors driving division plane selection in the root.

Plant cells undergo a variety of local and tissue-derived stress that may alter or influence division planes more directly. Wound-adjacent cells respond to their neighbours’ ablation through expression of stress-related genes like ETHYLEN RESPONSE FACTOR 115 (ERF115) (Heyman *et al.*, 2016; Marhava *et al.*, 2019; Canher *et al.*, 2020) and altered expansion towards the wound (Hoermayer *et al.*, 2020). Cell expansion represents a major source of mechanical stress in plant cells due to their rigidity and high internal turgor pressure. In the shoot apical meristem, division planes and microtubules (MT) align parallel to maximal tensile stress (Hamant *et al.*, 2008; Louveaux *et al.*, 2016). MTs are cytoskeleton filaments that are oriented perpendicular to the main expansion axis in elongating tissues (Kropf, Bisgrove and Hable, 1998) and mark the future position of the division plane in proliferating tissues (Rasmussen, Humphries and Smith, 2011). *In vitro* MT growth assays have shown that their stability is altered by stretching (Franck *et al.*, 2007; Trushko, Schäffer and Howard, 2013; Kabir *et al.*, 2014), suggesting that they may act as tensile stress sensors (Hamant *et al.*, 2019). However, *in vivo* demonstration for such activity is lacking and the possible mechanism linking tissue elasticity, cell expansion and cell division plane switches during tissue development remains elusive.

In this study we used cell ablation coupled with high-resolution and real-time imaging to study division plane switches during restorative cell divisions. We show that the whole tissue deforms, which is transmitted into reorientation of division planes in individual cells around the wound. This occurs via re-arrangement of MT arrays. Accordingly, MT stability is dependent on cell expansion and cell stretching, as inferred by wounding or through external aspiration, promotes MT bundle formation. Our results identify the mechanism, by which cell expansion, through applying stretching forces and thus reorienting MTs, orient division planes for tissue architecture.

## Results

### Directional expansion of cells correlates with division plane changes

Tissue regeneration in the root meristem involves induction of divisions and reorientation of division planes around the wound (Marhava *et al.*, 2019). To identify potential local and cell-cell communication mechanisms in response to wounds, we investigated the involvement of chemical signalling and found an accumulation of both cytosolic Calcium (Ca^2+^) and Reactive Oxygen Species (ROS; such as hydrogen peroxide H_2_O_2_). We observed the induction of Ca^2+^ waves starting instantly (within 0.2 s) from the ablated cells, transmitted through the neighbouring cells and reached the whole root tip within 25 s (Fig. S1A). Additionally, we observed an accumulation of H_2_O_2_, around the wound reaching a 1.22±0.12 fold increase within 18 min (Fig. S1B). However, the Ca^2+^ waves observed here did not correspond to the spatially restricted occurrence of restorative divisions. We also have previously shown that exogenous treatment of roots with H_2_O_2_ did neither induce wound responses like ERF115, nor trigger division plane changes (Hoermayer *et al.*, 2020). Additionally, when we applied an extract of wounded root tips to *ERF115::GFP* marker line seedlings, neither did we observe upregulation of ERF115 nor did the root extract induce any division plane changes (Fig. S1C). These observations did not support the idea of a causative chemical wound signal inducing restorative divisions.

Hence, we focused on a possibility that purely mechanical signals are involved and investigated the interdependence of cell expansion and restorative divisions by high-resolution 3D imaging. Notably, we observed division plane changes from anticlinal to periclinal in endodermis cells that were not in direct contact to the ablated cortex cells (Fig. 1A, white arrows). These cells increased in cell width compared to normal endodermis (green) cells in the same roots. Top view sections revealed that the width increased throughout the whole cell compared to neighbouring cells, marking an increase in cell volume (Fig. 1B, white arrows). Cells directly in contact with ablated cells increased similarly in width and volume (Fig. 1A and C, red arrows). To further investigate this phenomenon, we used long-term time-lapse imaging of wounded roots at the vertical stage microscope. This revealed that such non-adjacent cells undergo a constant cell expansion towards the wound (expansion rate was 1.44 times in 8.2 h before division, Fig. 1D-E). However, this expansion was significantly slower than the cell expansion observed for adjacent cells (1.47 times in 5h) (Fig. 1D-E). As control, we analysed the expansion of a neighbouring cell, which is more distant to the cell ablation than the non-adjacent cell. This neighbouring cell was expanding significantly less than the non-adjacent and adjacent cell (1.13 times in 9.8 h), and no alteration of the pre-established anticlinal cell division plane was observed (Fig. 1D-E, Video S1, Fig. S1F). This suggests that not the vicinity or connection to the wound induces division plane changes but expansion towards the wound.

**Figure 1.**
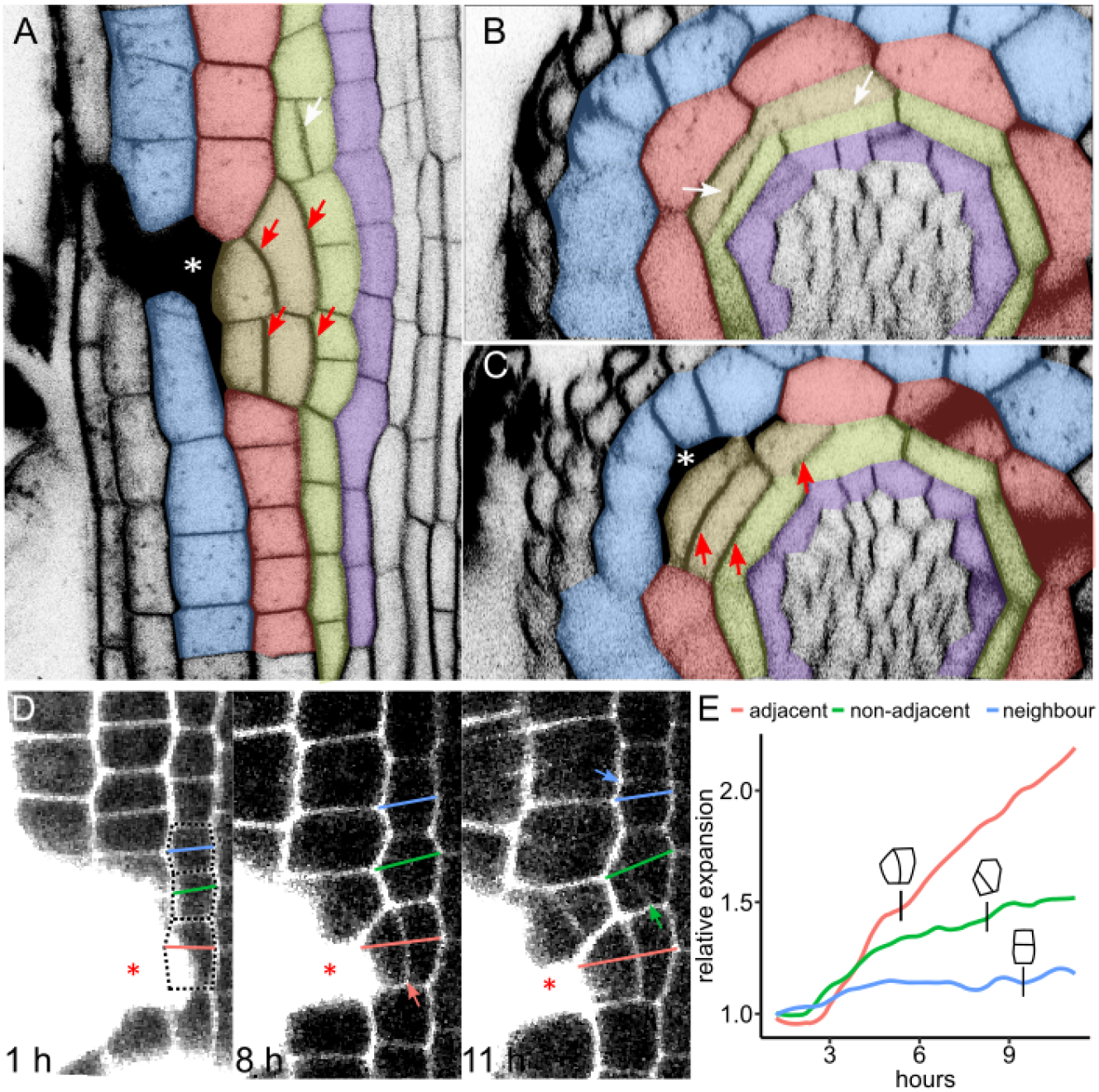
Division orientation defined by directional cell expansion. **(A-C)** Non-wound adjacent cells trigger division plane switch 24 hours after cortex and epidermis ablation. **(A)** Front view with periclinal divisions of wound-adjacent (red arrows) and non-adjacent (white arrows) endodermis. **(B)** Top view of non-adjacent endodermis cells (white arrows). **(C)** Top view showing the ablation with periclinal divisions in the endodermis (red arrows). Plasma membranes of root cells were stained with FM4-64. Cell types were re-colorized as epidermis (blue), cortex (red), endodermis (green) and pericycle (purple). **(D-E)** Non-adjacent periclinal divisions are preceded by cell expansion. **(D)** Time series of cell expansion close to ablation in propidium iodide (PI) stained root tip. Maximum cell width is drawn as line: red for adjacent cell, green for non-adjacent cell and blue for distant neighbour. Divisions are marked by arrows. **(E)** Quantification of cell width from (D) and time point of division. Graph is representative example (full quantification see Fig. S1F). Asterisks mark ablated cells.

Here we identified that the orientation of division planes after wounding can be predicted by occurrence of cell expansion towards the wound, which occurs not only by directly invading the wound area, but also by indirect stretching of non-adjacent cells.

### Wounding induces immediate outwards tissue deformation and cell expansion

The dependence of division plane changes on directional expansion or stretching motivated us to investigate the changes in mechanical properties of root tissues after wounding.

We have previously shown that the cell ablation causes an instantaneous pressure loss at the wound site (Hoermayer *et al.*, 2020). By time-lapse imaging, we found that the surrounding cells are rapidly pulled into the wounds within 20 min (Fig. 2A). Cells that were not in contact with the wound were also pulled towards the wound quickly (Fig. 2B, Video S2) suggesting that pressure changes immediately cause tissue deformation. Surprisingly, the cell displacement was not homogenous, and only the inner cells (with respect to the ablated cell) were pulled into the wound, while outer cells stayed mostly in place (Fig. 2A-C). To further dissect this phenomenon, we ablated two cell types simultaneously, the outer epidermis and inner endodermis with intact cortex cells in between. Both ablated cell types were replaced by the first respective inner adjacent cells (Fig. S2A-D). This result confirms that the inner cells respond more importantly in response to cell ablation.

**Figure 2.**
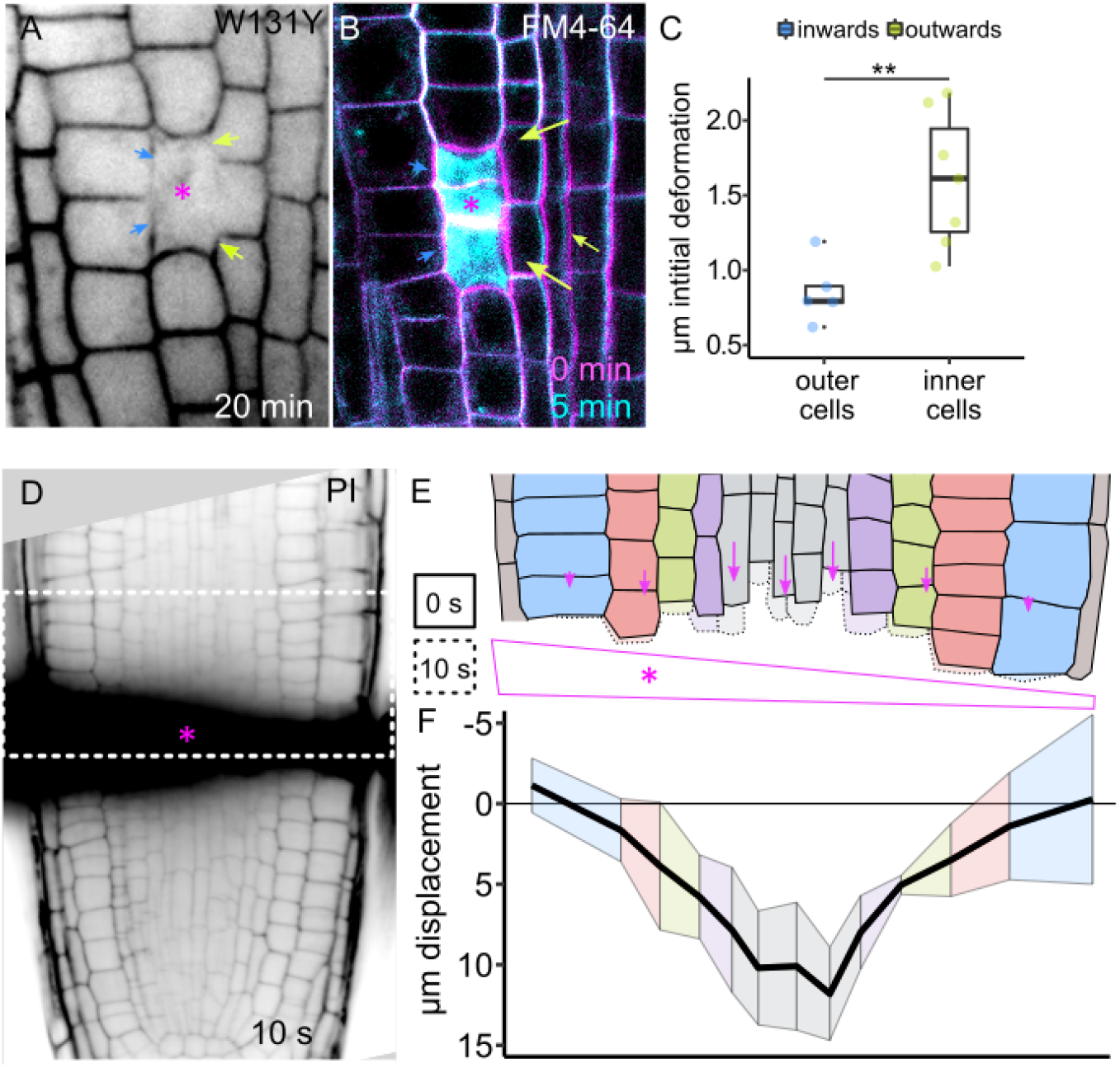
Instant, wound-induced deformations of root cells. **(A-C)** Root cell deformations induced after wounding. **(A)** *W131Y* plasma membrane marker 20 min after ablation. **(B)** Time series overlay of cortex cell ablation directly before death (magenta) and 5 min after death (cyan). Plasma membranes were stained with FM4-64. Arrows indicate displacement of membranes from inner (green) and outer (blue) cells. **(E)** Quantification of displacement in μm within first 20 min after ablation in outer cells inwards (blue) and in inner cells outwards (green). Boxplot marks median ± 95% confidence interval (CI). Statistical significance was computed from Wilcoxon test. **(D-F)** Extendibility of cells within the root meristem. **(D)** PI-stained root tip 10 s after ablation of a horizontal line of cells (black fluorescence). White dashed rectangular marks redrawn image in (E). **(E)** Redrawn section from (D) above horizontal ablation (marked as purple area with asterisk) before (full lines) and 10 s after ablation (dashed lines). Arrows indicate displacement. **(F)** Quantification of displacement towards wound in μm from (E). Black line indicates mean and coloured ribbon indicates 95% CI from n=3 roots. Cell types were colorized as epidermis (blue), cortex (red), endodermis (green), pericycle (purple) and stele (grey). Asterisks mark ablated cells.

To further investigate why inner cells deform and expand more towards wounds than outer cells, we ablated a horizontal section of cells, mimicking a macroscopic root tip excision (Fig. 2D). Intact cells around the ablation were immediately pulled into the wounded area (Fig. 2E, Fig. S2F, Video S3). The resulting cell wall displacement was strongest in cells of the stele, with a decreasing gradient towards the outer cell types (Fig. 2F). Notably, the displacement was transmitted also to cells further away from the ablation, reaching furthest in stele cells and decreasing in outer cells (Fig. S2E-F). This suggests that the root tissue behaves like an extendable matrix, with increased capacity to expand in the inner tissues.

Here, we revealed a tissue extendibility gradient in the root of more stretching inner cells to more rigid outer cells. This differential stretching within the root explains the preferential response of inner tissues mediating desired restorative division in response to wounding.

### Division plane is determined by a transient reorientation of microtubule arrays

Our results suggest that the induction of restorative divisions and the inherent division plane switch depend on cell expansion and their underlying physical deformations. However, the cellular and molecular machineries behind the division plane switch remain elusive. To address that, we first investigated the possibility of switching the division plane orientation throughout the cell cycle, mainly the M-phase where cytokinesis occurs and the directly preceding phase, G2.

We monitored the different cell division stages using the MTs marker line *35S::MAP4-GFP* (Marc *et al.*, 1998) and observed that adjacent cells already in mitosis were not able to switch their division plane (Fig. S3A-B). However, the spindles were highly tilted compared to control cells without ablations (Fig. S3A, Video S4). During cell division, the microtubule-rich pre-prophase band (PPB) marks the future cell division plane (Rasmussen, Humphries and Smith, 2011). Wound-adjacent cells with already established PPB at the moment of ablation divided without reorienting the cell division plane (Fig. S3C-D). However, using the G2-phase marker *CYCB1;1::GFP* (Ubeda-Tomás *et al.*, 2009) we observed wound-adjacent cells in G2 phase shortly before mitosis that still performed a division plane switch within 4.8-6.0 hours (Fig. S3E-F). These findings suggest that the orientation of division planes can still be switched late in the cell cycle, during transition to G2 phase. However, once the PPB is established and mitosis begins, the orientation of the cell division plane after ablation cannot be redefined.

Hence, we investigated the transition of cortical MT arrays to organized PPB rings. In the root meristem, cortical MTs visualized by the *35S::MAP4-GFP* marker line are oriented in weakly-defined transverse arrays (Fig. 3Ai-ii). Wound-adjacent cells underwent a re-arrangement of cortical MT orientation. Firstly, a new array of longitudinal cortical MTs appeared (Fig. 3Aiii), and secondly, a depletion of transversal MT arrays followed by the PPB formation in new orientation (periclinal) was observed (Fig. 3Aiv). Surprisingly, time-lapse imaging revealed that longitudinal arrays appeared when transversal arrays were still present and co-existed for additional 2.0 hours before the PPB formed (Fig. 3B). We observed single MT arrays during this co-existence phase and found multiple orientations of transversal, longitudinal and oblique bundles within the same cell (Fig. S3G). We used the CSBDeep deconvolution algorithm (Weigert *et al.*, 2018) for better visibility of MT arrays (Fig. 3C). Due to the small size of root cells and their stacked arrangement, a reliable quantification of cortical orientations was technically not possible, hence we measured the MAP4-GFP signal intensity on the membranes instead. The ratio of longitudinal versus transversal MT intensity was significantly higher in wound-adjacent cells (0.88 ± 0.122) compared to control cells (0.43 ± 0.147) (Fig. 3D, left).

**Figure 3.**
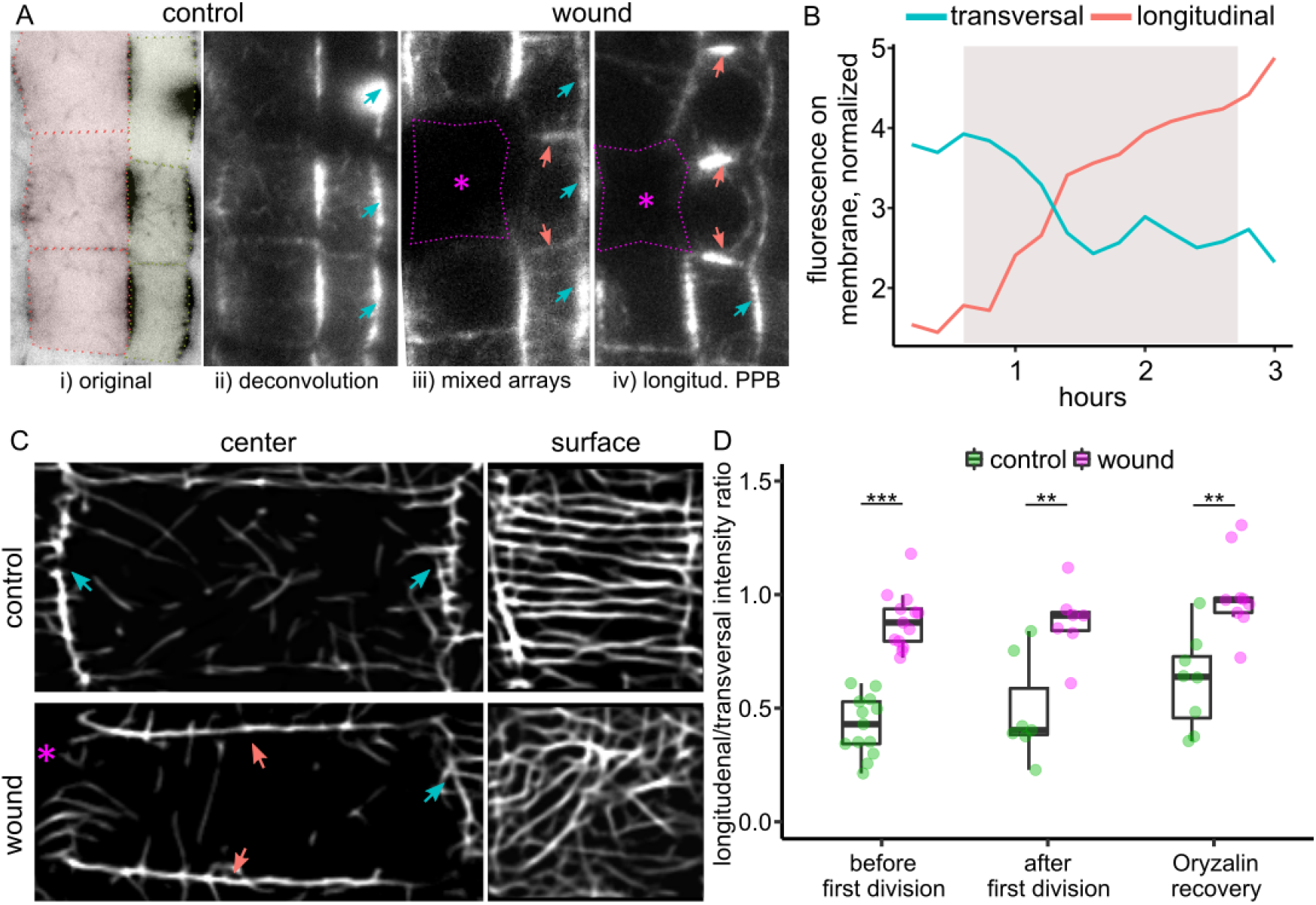
Wound-induced transient reorientation of microtubule arrays during restorative cell divisions. **(A-B)** Cortical MT array changes during restorative divisions. **(A)** From left to right: (i) Transverse MT arrays as marked by *35S::MAP4-GFP* in undisturbed cortex (red) and endodermis (green) cells. (ii) Inverted picture of (i) without cell type colorizing. (iii) Wound-adjacent endodermis cell with longitudinal and transverse cortical MT arrangements. (iv) Wound-adjacent endodermis cell with longitudinal-localized PPB and neighbouring cell with transverse arrays. Red and blue arrows indicate transversal and longitudinal arrays, respectively. Asterisks mark ablation. **(B)** Time-series quantification of *35S::MAP4-GFP* fluorescence at transversal/lateral (blue line) and longitudinal/apical (red line) membranes over time from one representative ablation site. Coloured area indicates co-existence of transversal and longitudinal arrays. **(C-D)** Multiple orientations of cortical-localized MT arrays. **(C)** Deconvolution of MT arrays as marked by *35S::MAP4-GFP* in control cortex cell (upper panels) and wound-adjacent cortex cell (lower panels). Left panels are from centre of cell and right panels are from cell surface with visible cortical MT bundles. Red and blue arrows indicate transversal and longitudinal arrays, respectively. Asterisks mark ablation. **(D)** Quantification of *35S::MAP4-GFP* ratio between longitudinal (apical membrane) and transversal (lateral membrane) fluorescence; (from left to right) before division, after division and during the recovery of the depolymerisation induced by Oryzalin 5 μM (12 hours). Magenta indicates wound-adjacent cells and green indicates control cells from the same roots. Statistical significance was computed using Wilcoxon test. Boxplot marks median ± 95% CI.

To investigate the origin of these multiple MT array orientations, we observed wound-adjacent cells after MT depolymerisation events. After the first division, where mitosis causes total depletion of MTs in the cell, the wound-adjacent daughter cells still displayed multiple MT orientations (Fig. 3D middle, Fig S3H). When we exogenously depolymerized MT cortical arrays using the depolymerisation drug Oryzalin (Morejohn and Fosket, 1991) and let the MT arrays re-polymerize after washout, we also observed multiple cortical MTs orientations in wound-adjacent cells (Fig. 3D right, Fig. S3I, Video S5). Notably, Oryzalin treated roots displayed normal wound-responsive expansion (Fig. S3J). Additionally, MTs oriented independently of actin polymerisation status (Fig. S3K).

These findings revealed that wound-adjacent cells switch MT orientation transiently from transversal to a mixed population of transversal and longitudinal before a complete division plane switch. These multiple orientations occurred independently of depolymerisation events.

### Cell expansion drives microtubule polymerization

The multiple orientations of cortical MT arrays in wound-adjacent cells could be a result of multiple directions of cell expansion that can include expansion towards the wound or following the growth axis of the root. To investigate this, we perturbed cell expansion in the root meristem and observed the MT array distribution in response.

Auxin inhibits cell expansion in the root elongation and transition zones (Fendrych *et al.*, 2018; Montesinos *et al.*, 2020). We observed similar inhibition of cell elongation in the meristem zone (Fig. S4A-C). This inhibited cell expansion was accompanied by a strong reduction of cortical MT arrays and increase of cytosolic fluorescence signal in the *35S::MAP4-GFP* marker line and as control, also in the *UBQ10::Venus-TUA6* marker line (Fig. S4D-E). To manipulate cell expansion by independent means, we treated roots with cellulose-biosynthesis inhibitor isoxaben (IXB) (Scheible *et al.*, 2001), which also reduced cellular growth in the root tip (Fig. S4F). Again, we observed a depletion of MT arrays during IXB treatment within 12 hours (Fig. S4G-H). Notably, cells in the transition zone (TZ) started to swell after prolonged IXB treatment, which was accompanied by an increased MT array formation (Fig. S4G, right panel). Our results suggest an interdependency of cell expansion and MT stability. Hence, we investigated what happens to MT arrays when cell expansion is rapidly induced.

We first inhibited cell elongation by 12 hours auxin treatment and performed single cell ablations to induce deformations. Any remaining transversal MT arrays completely disappeared within 1 min after ablation (Fig. S4I, left and middle panel) but notably re-polymerized within 15 min during the manifestation of the tissue deformation (Fig. S4I, right panel). To further dissect this observation, we performed laser ablations with weaker pulses, where we induced a delayed cell death. The harmed cell collapsed 5-10 min after ablation, and the wound-adjacent cells quickly deformed and stretched towards the wound (Fig. 4A middle panel). Notably, MT bundles in these wound-adjacent cells reappeared at the very moment of deformation (Fig. 4A middle panel). The *de-novo* induced bundles moved from the cell interior towards the cortical membranes within 30 min (Fig. 4A right panel, Fig. S4J, Video S6). Such an induction of MT bundles after neighbouring cell collapse was specific to directly adjacent cells (Fig. 4B), suggesting that MT bundles are directly initialised and stabilised by cell stretching.

**Figure 4.**
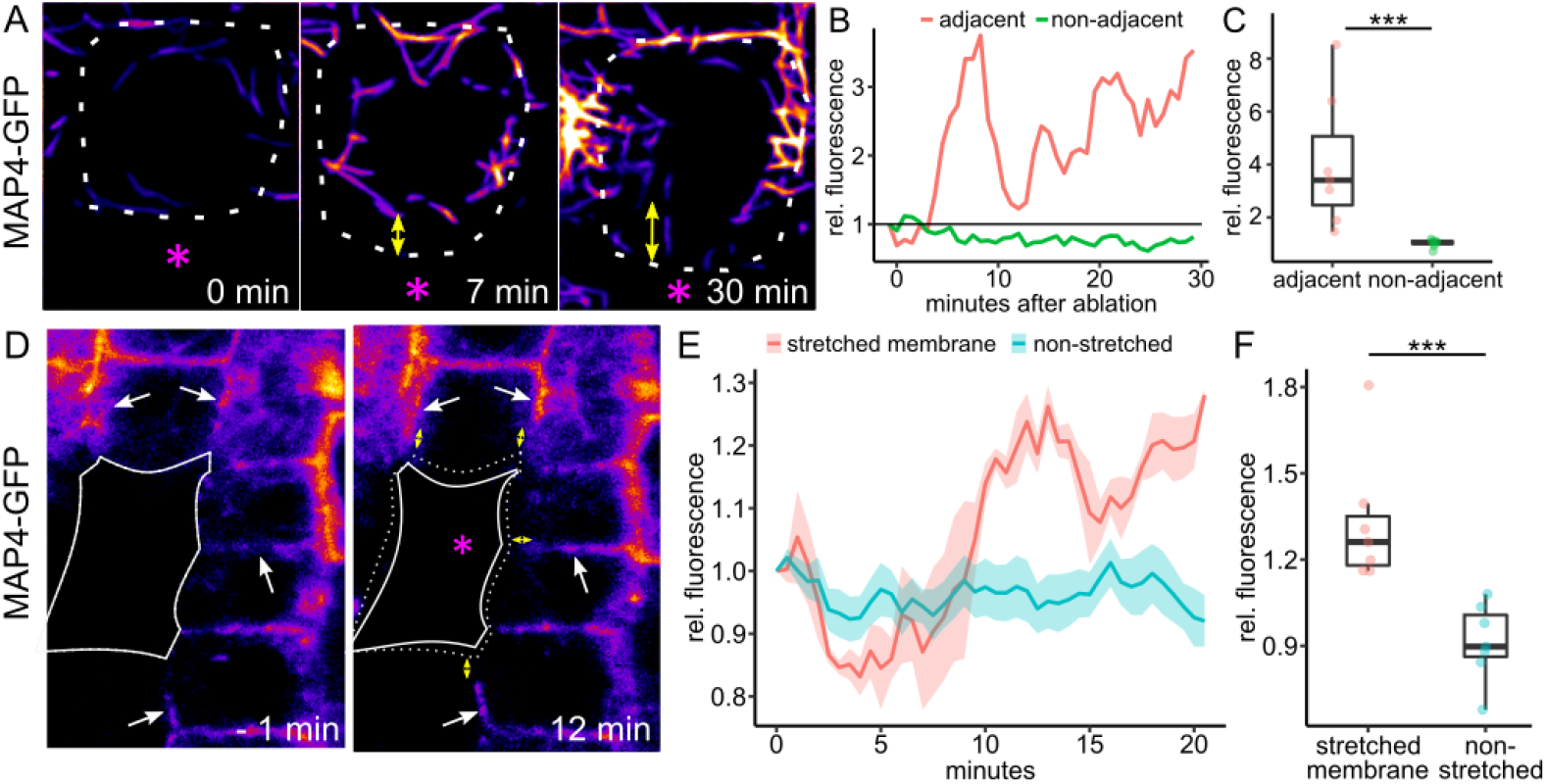
Induction of cell-localized MT polymerisation by cell stretching. **(A-C)** MT polymerisation is induced after the collapse of the neighbour cell. **(A)** Deconvolution images of MTs in a cortex cell adjacent to UV-laser harmed neighbour (asterisk. MT signal (*35S::MAP4-GFP*) before (left), during (middle) and after collapse of neighbouring cell (right). Roots were pre-treated with 1 μM NAA for 12 hours. Dotted line indicates stretching cell. **(B)** Time series of cytosolic *35S::MAP4-GFP* signal (mean grey value relative to t0) of a representative cell during stretching (red line) caused by collapse of an ablated neighbour and the closest neighbour non-adjacent to the collapsed cell (green line). **(C)** Quantification of the cytosolic *35S::MAP4-GFP* signal (mean grey value relative to t_0_) directly after cell collapse in adjacent cells undergoing stretching (red dots) and in non-adjacent cells (green dots). Statistical significance was computed using Wilcoxon test. Boxplot marks median ± 95% CI. **(D-F)** MT stability is enhanced on stretched membranes. **(D)** *35S::MAP4-GFP* marked MT arrays 1 min before (left) and after 12 minutes after (right) collapse of ablated cell. Yellow arrows mark stretched membranes and white arrows mark increased fluorescence at those membranes. **(E)** Time series of membrane-localised *35S::MAP4-GFP* signal (mean grey value relative to t_0_) from 3 cells neighbouring the ablation-induced cell collapse represented in (D). Stretched membrane signal (red line) and non-stretched membrane signal (blue line) have been taken from the same cells. Lighter background indicates standard error. (F) Quantification of the cortical *35S::MAP4-GFP* signal (mean grey value relative to t_0_) 12 min after cell collapse in wound-adjacent cells undergoing stretching (red dots) and not undergoing stretching (blue dots). Statistical significance was computed using Wilcoxon test. Boxplot marks median ± 95% CI.

In a complimentary experiment, we observed cortical MT abundance in cells around the wound before and after the cell collapse. Again, neighbouring cells were immediately stretched upon collapse of the ablated cell and cell sides perpendicular to the wound area increased slightly in length (Fig. 4D, yellow arrows). These stretched cell sides showed a significant increase of cortical MT signal in the *35S::MAP4-GFP* marker line (Fig. 4D, white arrows, Fig. 4E) 12 min after collapse of the ablated cell. Non-stretched membranes of the same cells did not show any increase in cortical MT abundance (Fig. 4E-F), suggesting a specific effect of collapse-induced stretching on cortical MT abundance.

Here, we revealed a direct effect of cell expansion, in the form of stretching after cell collapse, on MT stability. Our results indicate that stretching increases MT stability, causing an increased abundance of cortical MTs on stretched cell sides.

### Mechanical forces of externa stretching rapidly drive microtubule polymerization

To investigate more specifically whether MT polymerization is a response to mechanical forces of stretching, we adapted the micromanipulation-assisted micropipette aspiration system (Maître *et al.*, 2012) to plants and induced stretching of cells in living root tissue. To decrease the cell wall stiffness and turgor pressure we employed a mix of Macerozyme and Mannitol solutions, which lead to MTs depolymerisation preventing cortical arrays within roots (Fig. 5A, left panel). Notably, aspiration of epidermis cells caused an increase of GFP signal within the aspirated part of the cell, indicative of MT polymerisation events (Fig. 5A, middle and right panel). The MT signal increased by 0.06/min for 20 min and saturated to 2.3±0.59 times of pre-aspiration levels after 30 min (Fig. 5B, Fig. S5G). The aspirated cells displayed a 2.0±0.38 fold asymmetry of fluorescence signal between the aspirated part and the non-affected basal part within 30 min (Fig. 5C right panel, Fig. 5E). To discard possible unspecific increases of GFP signal induced by the stretching, we used *UBQ10::Venus-TUA6*, which showed the same effect and actin marker Fimbrin-YFP and *35S::GFP* (free cytosolic GFP), both showing no increase of fluorescence signal upon aspiration, respectively (Fig. S5A-G). To test, whether the increased fluorescence came from an increase of MT polymerisation, we treated plants with Oryzalin as depolymerisation drug and aspirated epidermis cells, resulting in no increase of fluorescence over time and no asymmetry between aspirated and non-aspirated parts (Fig. 5C, Fig. S5A, S5G). This indicates that the observed effect is specific to the formation of MT bundles and supports our previous findings that stretching increases MT stability.

**Figure 5.**
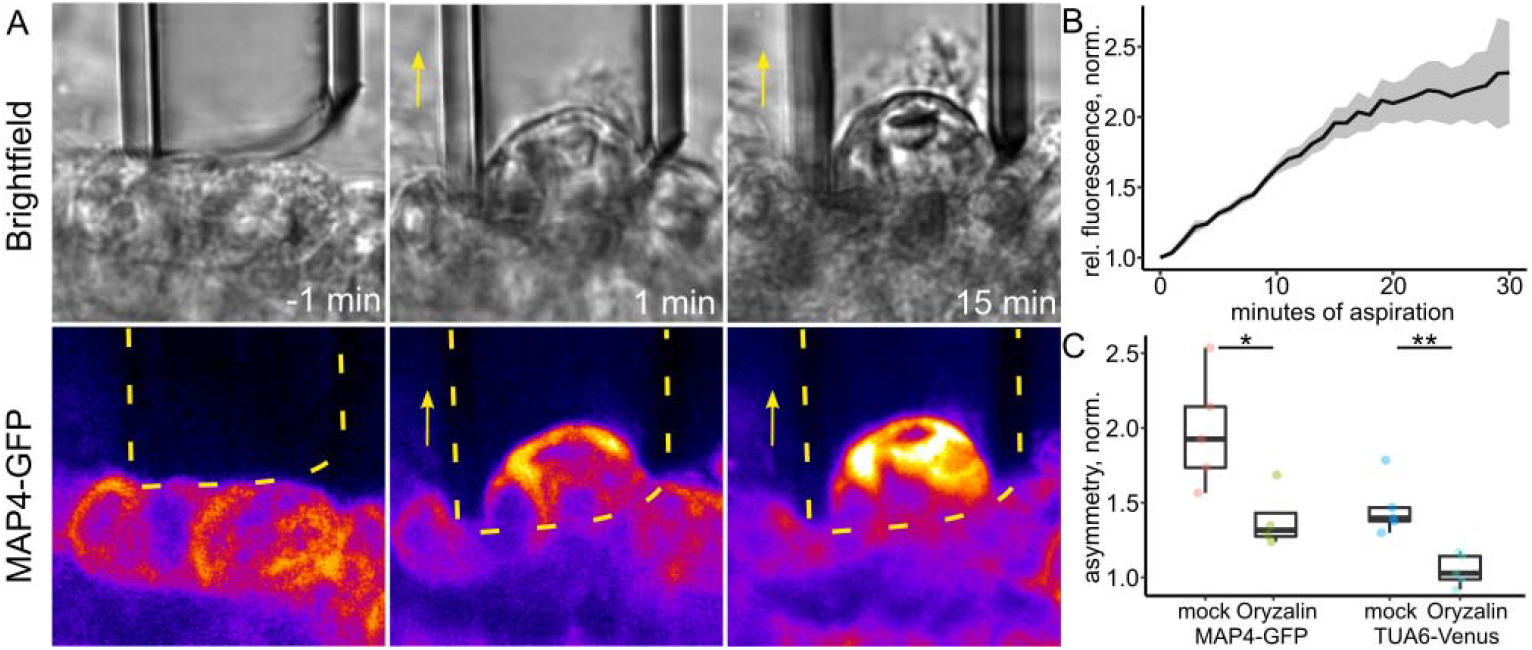
Induction of cell-localized MT polymerisation by aspiration. **(A)** Aspiration of epidermis cells. Top row: brightfield; bottom row: *35S::MAP4-GFP* fluorescence before (left), at the beginning (middle) and after 15 min (right) of aspiration. Yellow arrow marks direction of aspiration and yellow dotted line indicates position of pipette. **(B)** Time series of *35S::MAP4-GFP* signal in aspirated epidermis cells. Data are represented as mean from individual cells over time (mean grey value relative to t_0_, normalized by direct neighbouring cell) and lighter background indicates standard error. **(C)** Quantification of *35S::MAP4-GFP* (left) and *UBQ10::Venus-TUA6* (right) fluorescence of aspirated parts relative to non-aspirated part (=asymmetry) of epidermis cells during mock and 5 μM Oryzalin treatment. Data are normalized to closest non-aspirated neighbour. Statistical significance was computed using Wilcoxon test. Boxplot marks median ± 95% CI.

Thus, externally applied, micropipette aspiration-assisted cell stretching leads to immediate local MT stabilization supporting a direct link between mechanical forces and MT polymerization.

### Mechanical forces alter MT stability and drive division plane changes

Our experiments suggest that mechanical stress from stretching directly cause MT stabilization. This suggests that deformation and cell stretching after wounding predictably alter MT stability and orientation, subsequently driving division plane changes.

To visualize these effects, we quantified stretching and MT abundance changes over time from the same ablation site and represented them in a single, schematic image (Fig. 6). The ablation of epidermis cells was followed by an immediate deformation of the neighbouring cells due to the negative pressure from the collapsed cell. The negative pressure also changed mechanical forces within the root, leading to increased cell wall extension of up to 2-fold within 12 hours in transversal walls perpendicular to the wound area. The stretched transversal membranes show the strongest increase of cortical MT abundance over time (up to 1.2-fold), while non-stretched membranes remained unaffected and lost MT abundance over time (up to 0.5 fold). The increased abundance at stretched, transversal membranes lead to the occurrence of multidirectional MT arrays, with longitudinal/transversal MT ratios of 1.17 up to 1.43 in fully stretched cells, while non-stretched cells displayed ratios of 0.87 and below. Cells with longitudinal/transversal ratios of above 1.0 changed division planes from anticlinal (transversal) to periclinal (longitudinal), while cells with ratios below 1.0 remained the naturally occurring, anticlinal (transversal) division planes. These results show the direct connection of mechanical forces, here manipulated by cell ablation, with MT stability and orientation impacting division plane selection.

**Figure 6.**
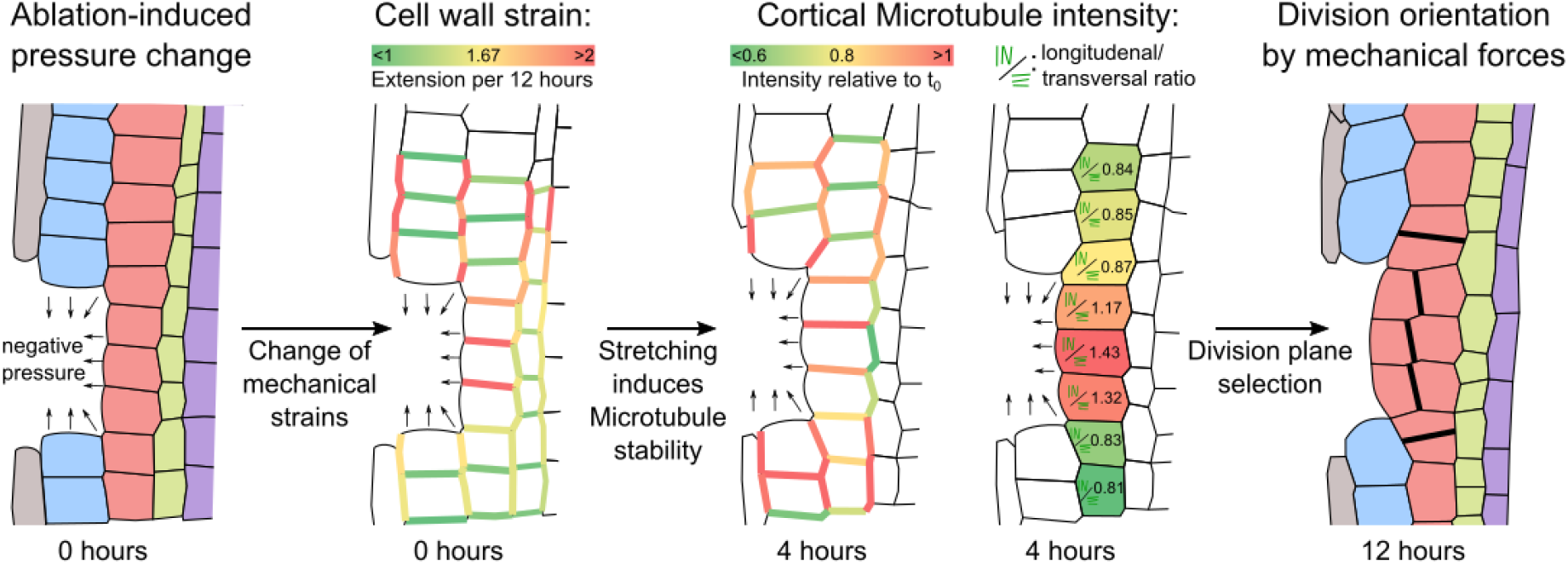
Schematic summary of mechanical effects after ablation on microtubule stability and orientation. Representative ablation of epidermis cells followed by expansion of cortex cells, changes in microtubule stability and orientation leading to changes in division plane orientation. From left to right: (i) Re-drawn and colorized confocal image of an epidermis ablation 10 minutes after ablation. (ii) Cell outlines and cell wall strain (as measured by relative cell wall extension after 12 hours) 10 minutes after ablation. (iii) Cell outlines and cortical MT intensity (measured as mean grey membrane signal relative to t0) 4 hours after ablation. (iv) Cell outlines and longitudinal/transversal MT array ratio (as measured by mean grey value on apical and basal relative to transversal membranes). (v) Re-drawn and colorized confocal image 12 hours after ablation showing new division planes as thick black lines. Cells were re-colorized as epidermis (blue), cortex (red), endodermis (green) and pericycle (purple). Small arrows indicate, negative pressure.

## Discussion

In this work, we focus on one of the key questions of developmental biology, namely, how plant cells switch their division planes – the key process, by which plant tissues and organs are shaped. We addressed this question in the context of restorative cell divisions during wound healing and tissue reconstruction in *Arabidopsis* root. We found that it is the direction of cellular growth and resulting mechanical forces, which directly affects cortical MT stability/polymerization leading to reorientation of MT arrays and ultimately new cell division planes. This data provides key insights towards understanding how cell division orientation can be flexibly rearranged in response to mechanical forces within the tissue and thus determine the plant architecture.

### Extracellular matrix transmits mechanical tensions for re-arrangement of cell growth

Plant cells are encapsulated by the cell wall, a cellular component that glues neighbouring cells together and prevents cell migration. Following wounding, adjacent cells and their neighbours further away are deformed and stretched towards the wound area. This shows that the extracellular matrix in roots is sufficiently plastic to transmit mechanical forces across the tissue. Stretched cells undergo division plane changes implying the importance of wounding-induced deformations for tissue restoration capacity. Notably, inner cell types are stretched more towards the wound than outer cells, suggesting a greater extendibility of inner tissues. Differential stiffness of certain cell walls in the root has been predicted (Fridman *et al.*, 2021), though experimental data has yet to support this idea (Elsayad *et al.*, 2016). Here, we observe for the first time differential extendibility in the root-cell wall matrix and show its relevance for developmental processes like wound healing and division plane determination, e.g. by assuring that wounds will be mainly healed from inside and not outside.

### Cell expansions strictly correlate with MT orientations and directions of cell divisions

The high turgor pressure of plant cells requires strong cell walls, which limit cellular expansion to occur in only one pre-defined direction. An ablation causes a pressure drop at the wound site that allows alternative directions of cell expansion, towards the wound. The changed expansion of wound-adjacent cells is accompanied by a re-arrangement of cortical MT arrays in two different directions: perpendicular to the main growth axis of the root (as originally in the undisturbed situation) and perpendicular to the direction of altered cell expansion towards the wound. Notably, these orientations persist also after depolymerisation and re-polymerisation of MTs. This supports the idea that cell expansion provides a continuous mechanical cue, e.g. through extension and re-arrangements in the cell wall itself (Kropf, Bisgrove and Hable, 1998; Fruleux, Verger and Boudaoud, 2019). Hence, division planes in the root meristem align perpendicular to the direction of maximum growth as can be seen in undisturbed situation, and pressure changes through wounding or other more physiological events can alter division plane orientation by physical effects alone.

### Cellular tensions are sufficient to rapidly drive MT polymerization

Wounding causes a local pressure drop that deforms the surrounding tissue and leads to continuous cell expansion towards a new direction. The deformation causes fast stretching of cell sides towards the wound. The immediate *de novo* MT polymerisation and subsequently increased abundance of cortical MT arrays at stretched sides suggests a direct link of mechanical forces of stretching with MT nucleation and stability. *In vitro* studies have suggested a connection of mechanical forces to MT polymerisation (Franck *et al.*, 2007; Trushko, Schäffer and Howard, 2013; Kabir *et al.*, 2014; Hamant *et al.*, 2019), however, no evidence for any *in vivo* effects have been shown so far. Using a novel approach of plant-cell aspiration, we show that induced tension is sufficient to stabilize MT polymerisation leading to formation of MT bundles. The purely mechanical stimulation causes MT polymerisation in stretched parts of the cell whereas non-aspirated parts remain unaffected. Similarly, after cell collapse, stretched cell sides show increased MT stability, while non-stretched sides remained unaffected. Over longer time, the constant stretching, changes MT array distribution and eventually results in MT reorientation. Hence, our data from internally and externally manipulated root cells provides evidence for a regulation of MT array formation by a directional cell stretching, thus constituting a mechanism of how cell growth via MT stabilization can rearrange MT orientation, which then ultimately determines cell division orientation.

## Conclusion

We identified a mechanism linking directional cell growth and orientation of the cell division, which depends exclusively on mechanical forces. These forces can be transmitted by an extendable extracellular matrix with differential rigidity leading to cell stretching, which is sensed through variations in MT stability/polymerisation allowing MT reorientation to ultimately determine the proper cell division plane. We demonstrated and visualised an interdependence of mechanical forces and MT polymerization *in vivo* and the underlying molecular mechanism linking both processes may or may not require other cellular components, i.e. cell wall receptors or MT associated proteins. Overall, these findings show how the directional cell expansion, via MT stabilization and reorientation, rearrange cell divisions planes ultimately determining tissue and plant architecture.

## Acknowledgements

We are thankful to Simon Gilroy, Alexander Jones and Lieven De Veylder for sharing published material. We thank the Bioimaging and Life Science Facilities at IST Austria for providing invaluable assistance. The research leading to these results has received funding from the European Research Council under the European Union’s Seventh Framework Programme (FP7/2007-2013) / ERC grant agreement n° 742985 and from the FWF under the stand-alone grant P29988.

## Author contributions

L.H., S.Y., P.M. and J.F. initiated the project. L.H., J.C.M., and J.F. designed experiments.

L.H., L.S. and S.C.-M. performed experiments and L.H. analyzed data. L.H., J.C.M. and J.F. wrote the manuscript. L.S., S.Y., P.M., S.C.-M., E.B. and C.-P.H. edited the manuscript. J.F. acquired funding.

## Declaration of interests

The authors declare no competing interests.

## List of Supplementary materials

Figure 1–6

Figure S1–S5

Methods

Videos S1-S6

## Supplementary Figures

**Figure S1.**
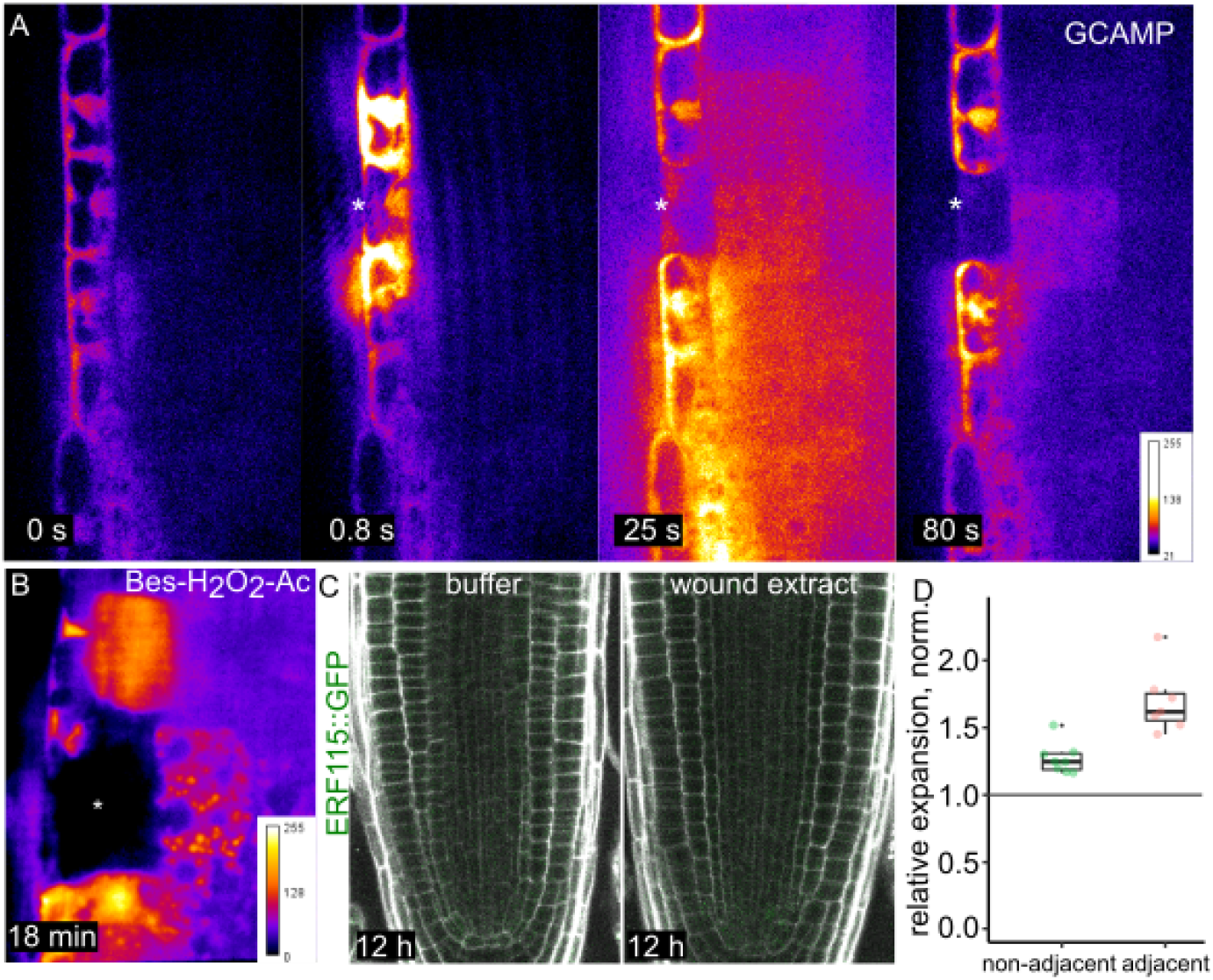
Chemical wound signals after ablation. **(A)** Calcium signal in root cells using the *GCaMP* calcium sensor during ablation of lateral root cap cell with time points at application of laser (0 s) and after 0.8 s, 25 s and 80 s (from left to right). Fluorescence intensity was obtained using GFP detection at a spinning disk microscope. **(B)** Intracellular H_2_O_2_ signal in root cells using the fluorescent dye BES-H_2_O_2_-Ac staining 18 min after ablation of epidermis cells. Note the strongest induction in wound-adjacent cells. **(C)** *ERF115::GFP* expression after 12 hours in phosphate buffer containing extract from wounded roots (right) or in mock conditions (buffer alone, left). Cell walls were stained with PI. Asterisks indicate ablated cells. **(D)** Relative cell width expansion towards ablated cells in non-adjacent cells (green) and adjacent cells (red) before periclinal divisions. Expansion was normalized to the closest neighbouring cell. Notice the significant increase of width expansion compared to neighbouring cells in both groups. Boxplot marks median ± 95% CI.

**Figure S2.**
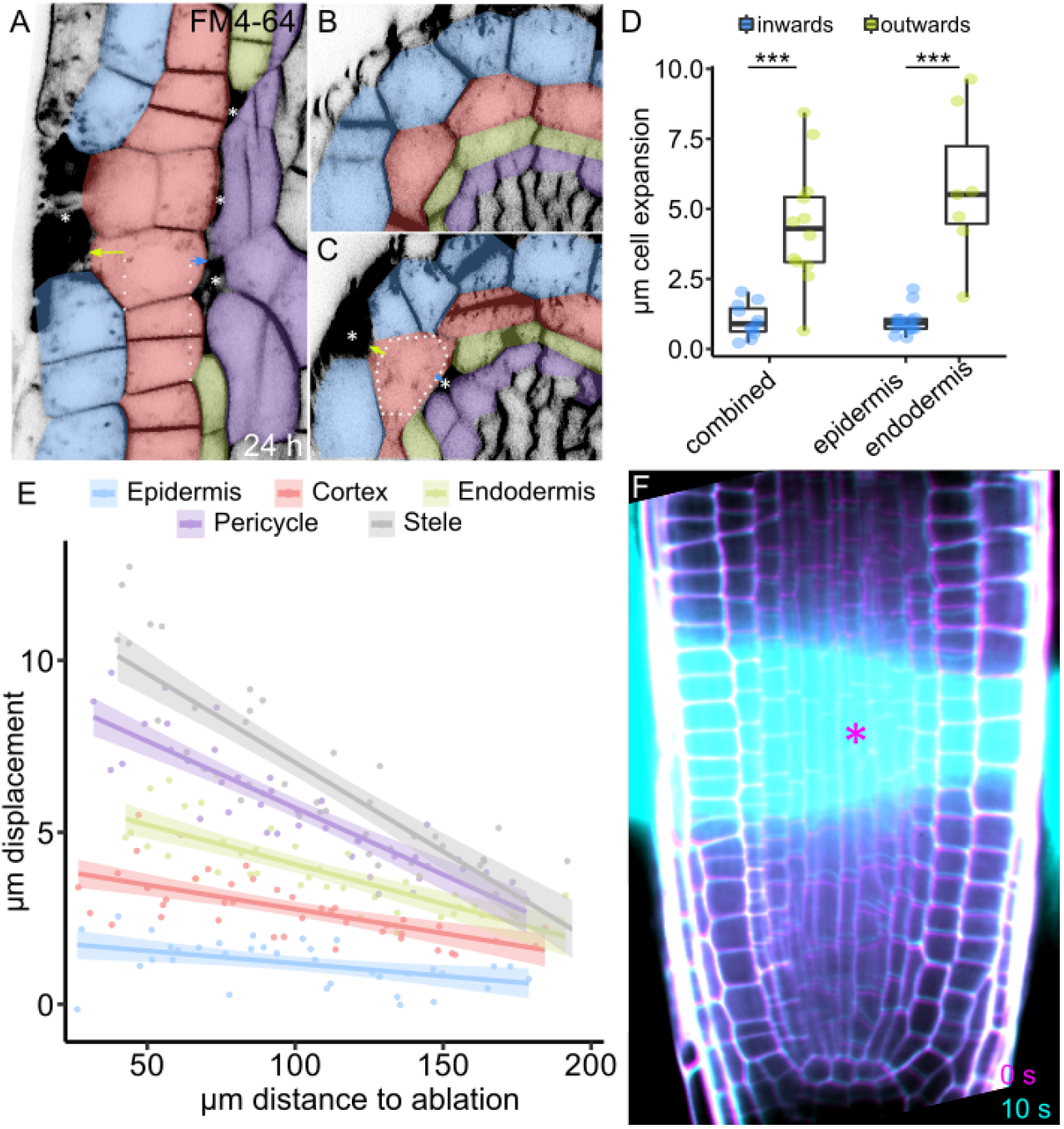
Wound-induced deformations. **(A-D)** Simultaneous ablation of epidermis and endodermis. **(A)** Front view with expansion of cortex cells outwards - towards ablated epidermis (blue arrow) - and inwards - towards ablated endodermis cells (green arrow). **(B)** Top view of different root cell layer in absence of cell ablation. **(C)** Top view directly adjacent to the ablation with expansion of cortex cells outwards - towards ablated epidermis (green arrow) - and inwards - towards ablated endodermis cells (blue arrow). Plasma membranes were stained with FM4-64. Cells were re-colorized as epidermis (blue), cortex (red), endodermis (green) and pericycle (purple). **(D)** Quantification of expansion of cortex cells 24 hours after ablation inwards (blue) and outwards (green) in combined ablations and separate ablations of epidermis and endodermis (from left to right). Boxplot marks median ± 95% CI. Statistical significance was computed from Wilcoxon test. Asterisks mark ablated cells. **(E-F)** Wound induced displacements are transmitted through whole tissue. **(E)** Quantification of displacement dependent on distance from ablation in μm. Dots represent single values, lines represent linear regressions and ribbons indicate 95% CI. **(F)** Time series overlay of horizontal section ablation before (magenta) and 10 s after ablation (cyan). Roots were stained with PI. Cell types were colorized as epidermis (blue), cortex (red), endodermis (green), pericycle (purple) and stele (grey).

**Figure S3.**
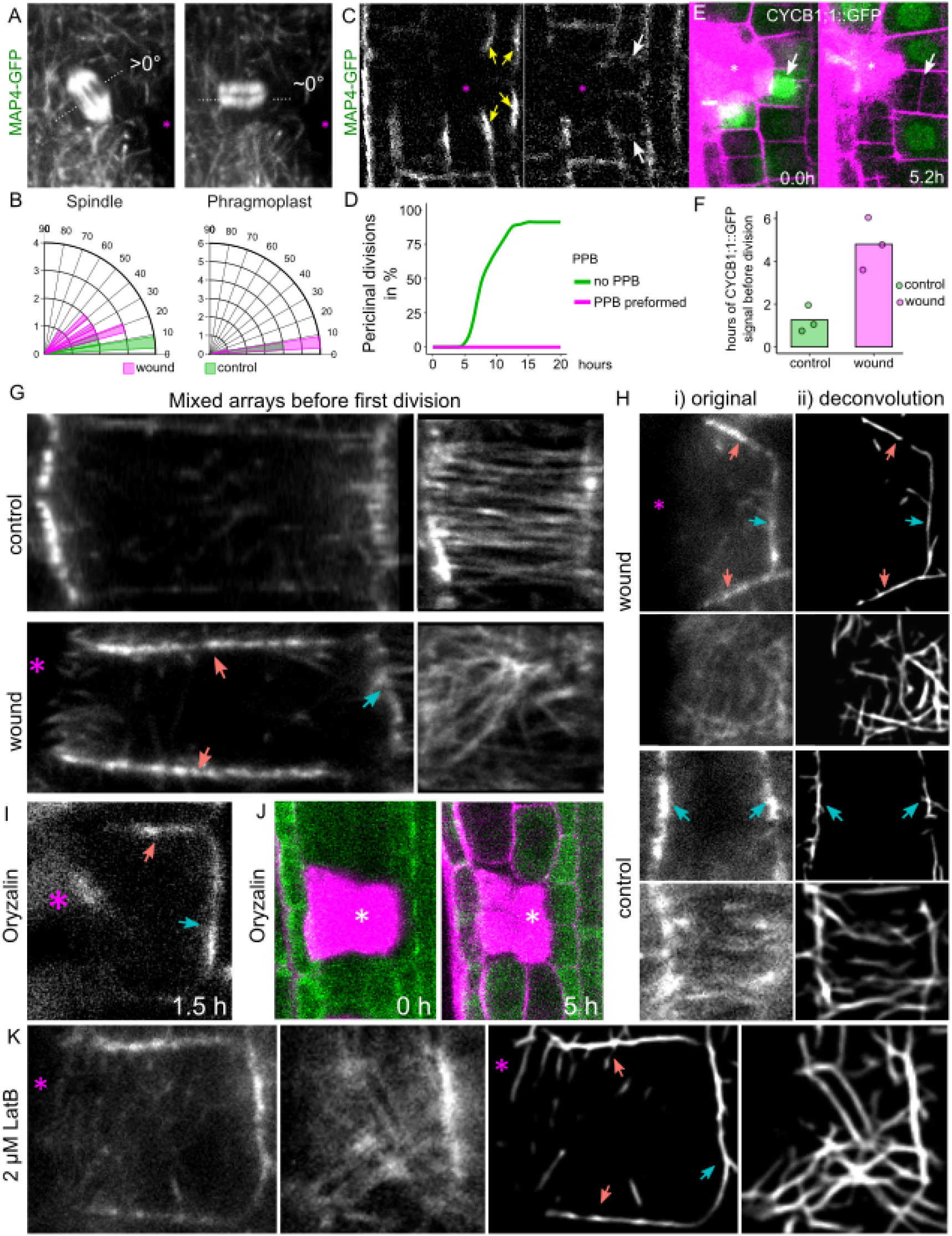
Microtubules during restorative divisions. **(A-B)** Cell division plane orientation during M-phase. **(A)** *35S::MAP4-GFP* marked spindle (left) and phragmoplast (right) after epidermis ablation (asterisk). **(B)** Quantification of angles of tilted spindle (left) and phragmoplast (right) in wound-adjacent (magenta) and distant (green) cells. Data are represented as histogram (bars). **(C-D)** Cell division plane during PPB formation. **(C)** *35S::MAP4-GFP* time-lapse analysis of wound-adjacent cells with pre-defined PPB (yellow arrows) at the moment of ablation (left) and after the first division (right). White arrows indicate anticlinal division. **(D)** Quantification of the rate of periclinal divisions over time in wound-adjacent cells with pre-defined PPBs (magenta) and in wound-adjacent cells without PPB at the moment of ablation (green). Data are represented as lines with data points every 0.2 hours. **(E-F)** Cell division plane change at onset of G2. **(E)** Time series experiment of wound-adjacent cells in G2 phase (as marked by green *CYCB1;1::GFP* expression) at the moment of ablation (left) and after the first division of the adjacent cells (right). Arrows indicate cells with CYCB1;1 expression that divided periclinally. Cells were stained with PI (magenta). **(F)** Quantification of the time between CYCB1;1 expression onset and division in wound-adjacent cells (magenta) and in distant cells (green). Asterisks mark ablated cells. **(G)** Original confocal images used for deconvolution images of Fig. 3C. **(H)** Cortical MTs (*35S::MAP4-GFP*) after first division. (i) Original confocal and (ii) deconvolution images. (“wound”) Cortical MTs in wound-adjacent endodermis daughter cells – cell centre (upper panels) and cell surface (lower panels). (“control”) Cortical MTs in non-adjacent endodermis daughter cells – cell centre (upper panels) and cell surface (lower panels). **(I)** Cortical MTs (*35S::MAP4-GFP*) in wound-adjacent cortex cell 1.5 hours after recovery from 12 h treatment with 5 μM Oryzalin. **(J)** Time series of root meristematic cells expressing the MTs marker *35S::MAP4-GFP* (green) upon 5 μM Oryzalin treatment during 5 hours after cell ablation (asterisk). Notice the predominately cytosolic localisation of the MTs signal. Cell walls were stained with PI (magenta). **(K)** Original (left panels) and Deconvolution images (right panels) of cortical MTs (*35S::MAP4-GFP*) in wound-adjacent cortex cells treated with 2 μM LatB 12 hours after ablation. Left: Cell centre, right: cell surface. Blue arrows: transversal arrays; red arrows: longitudinal arrays. Asterisks mark ablated cells.

**Figure S4.**
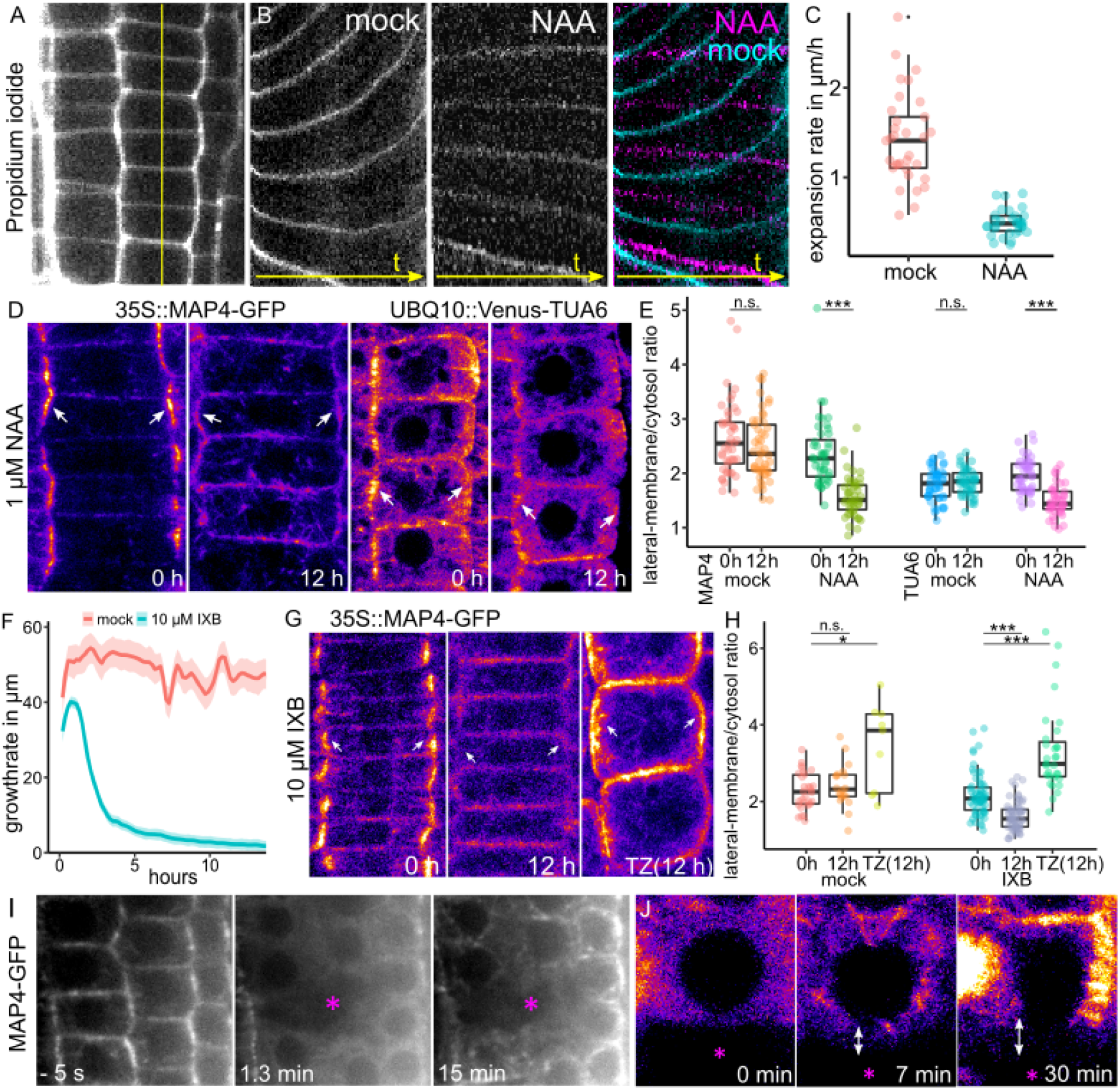
Auxin and Isoxaben effect on cell expansion and microtubules. **(A-B)** Auxin inhibition of MT array abundance. **(A)** *35S::MAP4-GFP* (left panels) and *UBQ10::Venus-TUA6* (right panels) marked MT arrays after 0 hours and after 12 hours of 1 μM NAA treatment. White arrows indicate lateral membrane intensity. **(B)** Quantification of *35S::MAP4-GFP* and *UBQ10::Venus-TUA6* fluorescence (lateral membrane relative to cytosolic signal) at 0 and 12 hours after mock (red and blue) and 1 μM NAA treatment (green and purple). Statistical significance was computed using Wilcoxon test. Boxplot marks median ± 95% CI. **(C-D)** Isoxaben (IXB) inhibition of MT array abundance. **(C)** *35S::MAP4-GFP* marked MT arrays after 0 hours (left) and after 12 hours (middle) of 10 μM IXB treatment in meristematic zone and after 12 hours (right) in transition zone (TZ). White arrows indicate lateral membrane intensity. **(D)** Quantification of *35S::MAP4-GFP* fluorescence (lateral membrane relative to cytosolic signal) at 0h and 12 hours in meristematic and transition zone (TZ) after mock (left) and 10 μM IXB treatment (right). Statistical significance was computed using Wilcoxon test. Boxplot marks median ± 95% CI. **(E-G)** Auxin inhibition of cell expansion. **(E)** Meristematic root cells stained with PI. Yellow line indicates slice of cortex cell walls used for generating kymograph (mock) of (G). **(F)** Kymographs of PI-stained cortex cell walls upon mock (left), 1 μM NAA treatment (middle) and overlay (right). Time is displayed on the x-axis, one pixel is 0.2 h and total time is 17.2 h. **(G)** Relative cell expansion rate (μm/h) of cortex cells during growth of 24 hours with mock (red) or 1 μM NAA (blue) treatment (from kymographs). Boxplot marks median ± 95% CI. **(H)** Isoxaben inhibition of root growth. Quantification of root growth during vertical stage microscopy. Lines represent smoothed mean of root growth in μm per 0.2 hours and lighter background indicates standard error. **(I)** Time-lapse confocal images of meristematic root cells expressing *35S::MAP4-GFP* (MT arrays marker) before (−5 s) and after (1.3 and 15 min) cell ablation in roots pre-treated with 1 μM NAA for 12 hours. Notice the loss of MT arrays immediately after laser application (middle) and the MTs re-polymerisation after the collapse of ablated cell (right). (J) Original confocal images used to obtain the deconvolution images of Fig. 4A. Asterisks mark ablated cells. White arrows mark cell stretching.

**Figure S5.**
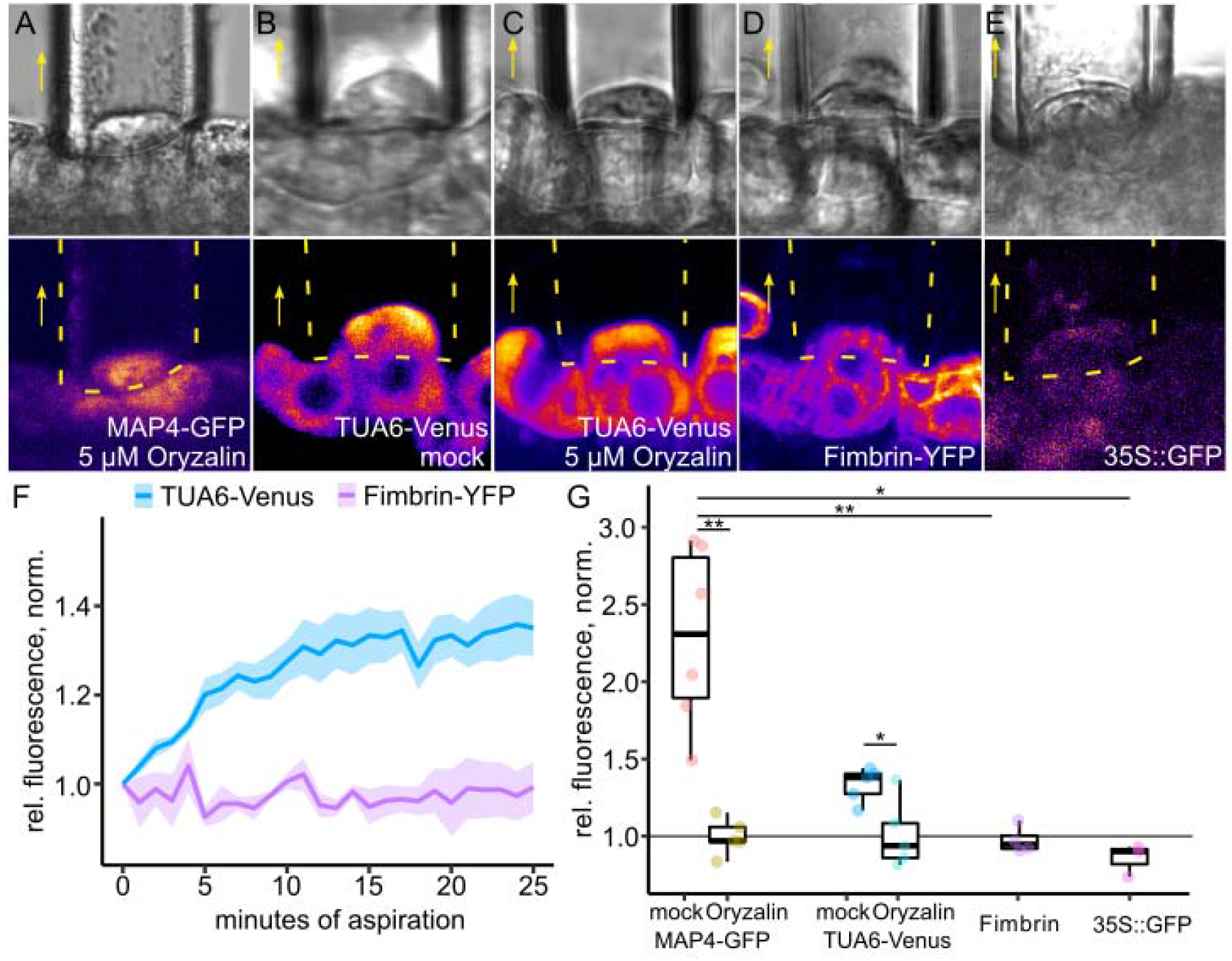
Dynamics of microtubules during aspiration assays. **(A-E)** Aspiration of epidermis cells 20 min after aspiration. **(A)** *35S::MAP4-GFP* upon 5 μM Oryzalin treatment. **(B)** *UBQ10::Venus-TUA6* upon mock treatment. **(C)** *UBQ10::Venus-TUA6* upon 5 μM Oryzalin treatment. **(D)** Actin filaments marked by *35S::Fimbrin-GFP.* **(E)** Free, cytosolic GFP marked by *35S::GFP.* Yellow arrows mark direction of aspiration. Yellow dotted lines indicate position of micropipette. **(F)** Time series of *UBQ10::Venus-TUA6* (blue) and *35S::Fimbrin-GFP* (purple) signal in aspirated epidermis cells. Data are represented as mean from individual cells over time (mean grey value relative to t0, normalized by direct neighbouring cell) and lighter background indicates standard error. **(G)** Quantification of fluorescence of aspirated epidermis cells during mock and 5μM Oryzalin treatment after 20 min of aspiration. Data are represented as mean grey value relative to t_0_ and normalized to closest non-aspirated neighbour. Statistical significance was computed using Wilcoxon test. Boxplot marks median ± 95% CI.

## METHODS

### Plant material

*Arabidopsis thaliana* (L.) Heynh (accession Columbia-0) was used in this work (WT). The following transgenic *Arabidopsis thaliana* lines were described previously: *35S::MAP4-GFP* (Marc *et al.*, 1998), *CYCB1;1::GFP* (Ubeda-Tomás *et al.*, 2009), “GCAMP” *35S::GCaMP3* (Toyota *et al.*, 2018), *35S::Fimbrin-GFP* (Wang *et al.*, 2004), *W131Y* (Geldner *et al.*, 2009), *ERF115::NLS-GFP-GUS* (Heyman *et al.*, 2016).

### Growth conditions

Seeds of *A. thaliana* were sown on Murashige and Skoog (1/2MS) medium (Duchefa) with 1% sucrose and 1% agar, stratified for 1-2 d and grown for 3-5 d at 21°C in a 16 h light/8 h dark cycle.

### Pharmacological treatments

Seedlings were transferred on solid MS medium containing the indicated chemicals: propidium iodide (PI, 10 μM, Sigma-Aldrich or Thermofisher), Naphthylacetic acid (NAA, Duchefa Biochemie, final concentration 1 μM), Mannitol (Sigma-Aldrich, final concentration 0.2 M), Isoxaben (Sigma Aldrich, final concentration 10 μM), Latrunculin B (LatB, Sigma-Aldrich, final concentration 2 μM), Oryzaline (Duchefa Biochemie, final concentration 5 μM), Macerozyme R-10 (Serva, final concentration 1% w/v).

### Sample preparation

Seedlings were placed on chambered cover glass (VWR, Kammerdeckgläser, Lab-Tek™, Nunc™ - eine kammer, catalog number: 734-2056) as described (Marhavý and Benková, 2015). With the chamber, a block of solid MS media was cut out and propidium iodide solution was added. After the liquid soaked in, 10-15 seedlings were transferred to the agar and the block was inserted into the chamber. For FM4-64 staining, seedlings were incubated in FM4-64 staining solution (3 μg/ml in water) for 3 min and then placed on solid MS medium.

### Confocal imaging and image processing

Confocal imaging was performed with Zeiss LSM700/800 inverted microscopes using 20x or 40x objectives. Detection of fluorescence signals was carried out for GFP (excitation 488 nm, emission 507 nm), YFP (excitation 514 nm, emission 527 nm) and PI or FM4-64 (excitation 536 nm, emission 617 nm). For fixed time point measurements, samples were observed 12 hours after ablation or at indicated time points. Images were analysed using the ImageJ (NIH; http://rsb.info.nih.gov/ij) and Zeiss Zen 2.3 “Black” or “Blue” software. Where necessary, images were processed by adjusting contrast and lightness. Where indicated, images where processed by sum slice operation to summarize multiple confocal planes or time points into one image. For 3D images, confocal (z) planes were chosen to fit ½ airy unit. To obtain top view images, z-stacks were processed using the “Reslice” tool.

### Spinning disk imaging

For the observation of immediate effects during/after ablation, an Andor spinning disk microscopy (CSU X-1, camera iXon 897 [back-thinned EMCCD], FRAPPA unit and motorized piezo stage) 63× water immersion objective was used. Videos were acquired with 1 focal plane, every 0.2 or 1.0 seconds. All images in a single experiment were captured with the same settings.

### Vertical stage microscopy, root tracking and image processing

Vertical stage microscopy for long-term tracking of root meristems was performed as described (von Wangenheim *et al.*, 2017; Glanc, Fendrych and Friml, 2018; Marhava *et al.*, 2019). Roots were imaged with a vertically positioned LSM700 or LSM800 inverted confocal microscope and Zeiss Zen 2.3 “Black” or “Blue” software, respectively, with 20x objective and detection of PI, GFP (see above) and transmitted light. Z-stacks of 30-42 μm were set accordingly to image each cell at least once. For the root-tracking, the TipTracker MATLAB script (Zen Black) or the TipTracker internal macro (Zen Blue) were used; interval duration was set between 600 s (10 min) and 720 s (12 min). The resulting images were concatenated and analysed using ImageJ. For registration, ImageJ macros “correct 3D drift”, “StackReg” or “MultiStackReg” were used. Kymographs were generated using the “Reslice” tool.

### UV laser ablation setup

The UV laser ablation was performed as described in ref. (Marhavý *et al.*, 2016; Marhava *et al.*, 2019) which are based on the layout published in ref. (Colombelli, Grill and Stelzer, 2004). The laser was applied in the upper corner of the cell of interest.

### Aspiration setup

The aspiration setup was designed as previously described (Maître *et al.*, 2012). 20 μm glass pipettes (BioMedical Instruments) with 30° bent angle were filled with distilled water and then connected to a Microfluidic Flow Control System (Fluigent, Fluiwell), with negative pressure ranging from 7−750 Pa. The microfluidic setup was mounted on a micromanipulator (Eppendorf, Transferman NK2) and micropipette movement and pressure were controlled via a custom-programmed Labview (National Instruments) interface.

5-d old seedlings were transferred onto 1% Macerozyme and 0.2 M Mannitol containing agar medium. After 20 min the seedlings were transferred carefully to a 0.2 M Mannitol containing agar medium to recover from the cell wall digestion. Finally, the seedlings were transferred to liquid medium containing 0.2 M Mannitol in a MatTek 50mm Glass Bottom Dish, so that the cotyledons touched the glass and the root was facing towards the centre of the coverslip.

For the aspiration, the micropipette was placed touching a single epidermis cell in the centre of the root meristem and during imaging, full negative pressure (−750 Pa) was applied. Imaging was performed using a Leica SP5 or a Leica Stellaris 5 confocal microscope with a resonant scanner and a Leica 20X, 0.7 NA objective, (Argon laser: 488 nm) for simultaneous imaging of fluorescent and brightfield channels.

### Wounded root extraction

Seeds were sown on a sieve placed in a jar filled with MS medium to cover the seeds. After 4 weeks the root tips were harmed with a scalpel, and after 10 min incubation, the shoot was cut off and the roots were ground with a mortar and constantly cooled with liquid nitrogen. The root extract was stored at −80°C. 5 ml of a phosphate buffer containing 38mM KH□PO□ and 62mM K□HPO□ were cooled to 4 °C and added to 1 g of the root extract. After an incubation time of 5 minutes, the suspension was centrifuged at 15000g for 20 minutes at 4 °C. The supernatant was applied to growing 5-day-old seedlings (1 ml per hour to each root continuously for 12 hours).

### 3D visualization and Deconvolution

3D images were obtained and analysed using Imaris software. Deconvolution images were generated by CSBDeep Fiji plugin – option “Deconvolution – Microtubules” with default settings.

### Quantification and statistical significance

Asterisks illustrate the p-value: p < 0.001 is ***, p < 0.01 is ** and p < 0.05 is * Boxplots represent median +/−95% confidence interval (= 1.58*IQR/sqrt(n))

### Spindle and phragmoplast angles

Spindle/phragmoplast angle was determined using the ImageJ “Angle” tool, taking the anticlinal walls as reference (0°). Data was plotted as histogram on a polar coordination system using R.

### Periclinal divisions

Division events were counted by marking the time point at which a new cell wall (stained by PI) appeared in the first, inner adjacent cell of the ablation site. Only periclinal divisions (vertical cell walls) were counted. The percentage of cumulative division events over time was plotted using R.

### Fluorescence intensity in single cells within time series

Signal intensity (mean grey value) from multi-stack (3D) videos in the green channel was quantified using ImageJ and recorded for each available time frame. To obtain relative values, the raw data was divided by the first time point value. Similarly, data from controls or reference cells (neighbouring or more distant cells) was recorded and the ratio of sample to reference value was calculated to obtain normalized values.

### Cell expansion

Cell expansion within time series was approximated by the cell width, quantified as the distance between the midpoint of the inner and outer cell wall using ImageJ. Expansion values are relative to first time point and if indicated, normalized to neighbouring (reference) cells.

During fixed time points, cell expansion or initial displacement was approximated as the horizontal distance between the wound-adjacent membrane and the first, non-adjacent membrane of a neighbouring cell (see dotted lines in Fig. 3A).

Expansion rates were determined from kymographs of growing root tips. One patch of cortex cells was registered over time, turned into a kymograph, coordinates of each line were obtained using Kymobutler(Jakobs, Dimitracopoulos and Franze, 2019), and the rate of expansion was calculated from the slope of the lines.

### Microtubule intensity within time series

Microtubule intensity (*35S::MAP4-GFP*) in growing root tips upon auxin treatment was measured in selections from one confocal plane over time, obtained from the “Threshold” tool in ImageJ. Every time frame was evaluated separately with a new selection. To obtain relative values, the raw data was divided by the first time point value.

Microtubule intensity (*35S::MAP4-GFP*) in growing root tips upon isoxaben treatment was measured in maximum z-projections over time in ImageJ. All time frames were evaluated with the same selection of the indicated third of the meristem. To obtain relative values, the raw data was divided by the first time point value.

Microtubule array intensity of a single cell was measured in a rectangular section from one confocal plane over time. All time frames were evaluated with the same selection.

Microtubule bundle intensity of a single cell during collapse of the neighbouring cell was measured in both, the first available time frame before collapse and after collapse and relative values were obtained by division. Values were normalized by division with values from neighbouring cells, not adjacent to collapsed cells.

### Microtubule intensity ratios

For lateral/apical membrane intensity ratio, microtubule intensity (*35S::MAP4-GFP*) was measured on the apical membrane and on the lateral membrane not touching the wound. Reference/control cells were chosen from the same cell type of the same root further away from the wound (no visual deformations).

For lateral-membrane/cytosol intensity ratio, microtubule intensity (*(35S::MAP4-GFP* and *UBQ10::Venus-TUA6*) was measured on the inner lateral membrane and in the cytosol excluding nucleus area.

### Microtubule intensity within aspiration assays

Microtubule intensity (*35S::MAP4-GFP*) in epidermis cells was measured in selections from one confocal plane over time. All time frames were evaluated with the same selection including reference cells (neighbouring, non-aspirated cells). Time series data was smoothed by taking the mean of a period of 1.8 s (5 frames for 0.37 s interval and 9 frames for 0.2 s interval). No smoothing has been applied for long-term aspiration (10 min interval). Relative values were obtained from division with first value and normalized values from division with reference cells.

## Video legends

**Video S1. Non-adjacent periclinal division are preceded by cell expansion.**

Cell expansion in wound-surrounding cells after epidermis and cortex ablation in propidium iodide stained *Arabidopsis* root meristem. Asterisk marks ablation. Green arrow indicates non-adjacent cells undergoing cell expansion and division plane switch. Quantification see Fig. 1E.

**Video S2. Instant, wound-induced deformations in root cells.**

Two consecutive ablations of one epidermis and one cortex cell in propidium iodide stained *Arabidopsis* root meristem.

**Video S3. Root cell extendibility after horizontal ablation.**

Full, horizontal-line ablation in propidium iodide stained *Arabidopsis* root meristem.

**Video S4. Spindle phragmoplast orientation in wound-adjacent cells**

*35S::MAP4-GFP* marked spindle and phragmoplast in root cell after ablation in the moment of nuclear breakdown.

**Video S5. MT orientation in wound-adjacent cells during re-polymerization**

Cortical MTs (*35S::MAP4-GFP*) in wound-adjacent cortex cell (white arrow) after recovery from 12 h treatment with 5 μM Oryzalin. Asterisk marks ablation. Red and blue arrows indicate longitudinal and transversal MT arrays, respectively.

**Video S6. MT re-polymerization is induced after the collapse of the neighbour cell.**

Visualization of MTs (*35S::MAP4-GFP*) in a cortex cell (white arrow) adjacent to UV-laser harmed neighbour (purple asterisk) after 12 hour 1 μM NAA pre-treatment.

## Notes

### Competing Interest Statement

The authors have declared no competing interest.

